# The Global Register of Introduced and Invasive Species: Country Compendium

**DOI:** 10.1101/2022.04.19.488841

**Authors:** Shyama Pagad, Stewart Bisset, Piero Genovesi, Quentin Groom, Tim Hirsch, Walter Jetz, Ajay Ranipeta, Dmitry Schigel, Yanina V. Sica, Melodie A. McGeoch

**Author notes:** Corresponding authors: Shyama Pagad; Melodie A. McGeoch.

## Abstract

The Country Compendium of the Global Register of Introduced and Invasive Species (GRIIS) is a collation of data across 196 individual country checklists of alien species, along with a designation of those species with evidence of impact at a country level. The Compendium provides a baseline for monitoring the distribution and invasion status of all major taxonomic groups, and can be used for the purpose of global analyses of Introduced (alien, non-native, exotic) and Invasive species (invasive alien species), including regional, single and multi-species taxon assessments and comparisons. It enables exploration of gaps and inferred absences of species for countries, and provides a means for short to medium term refinement of GRIIS Checklists. The Country Compendium is, for example, instrumental, along with data on first records of introduction, for assessing and reporting on invasive alien species targets, including for the Convention on Biological Diversity and Sustainable Development Goals. The GRIIS Country Compendium provides a baseline and mechanism for tracking the spread of introduced and invasive alien species across countries globally.

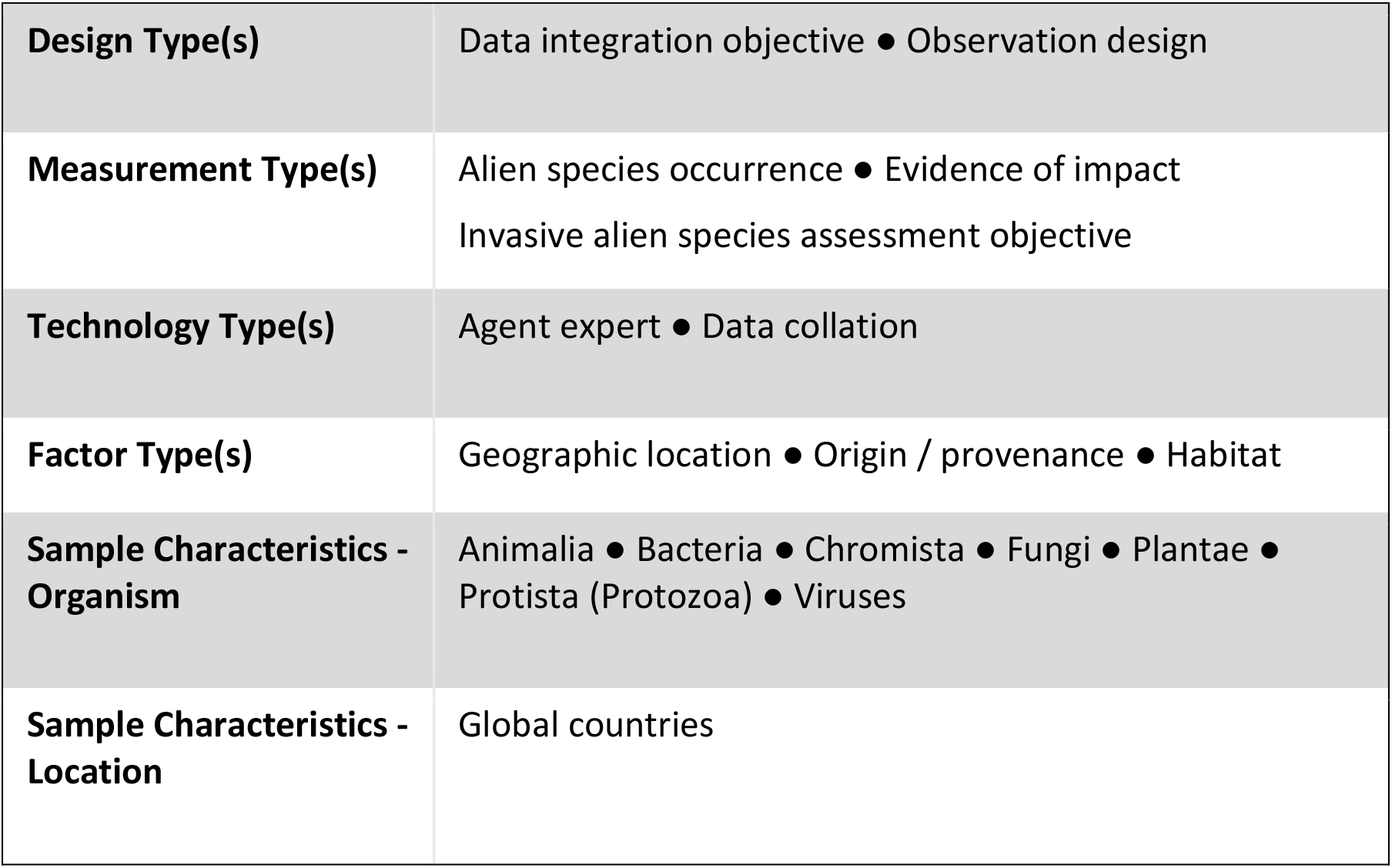

|  |  |
| --- | --- |
| **Design Type(s)** | Data integration objective • Observation design |
| **Measurement Type(s)** | Alien species occurrence • Evidence of impact<br>Invasive alien species assessment objective |
| **Technology Type(s)** | Agent expert • Data collation |
| **Factor Type(s)** | Geographic location • Origin / provenance • Habitat |
| **Sample Characteristics - Organism** | Animalia • Bacteria • Chromista • Fungi • Plantae • Protista (Protozoa) • Viruses |
| **Sample Characteristics - Location** | Global countries |

## Background & Summary

The need for up-to-date information on introduced species that harm the natural environment grows alongside their numbers, distributions and impacts on biodiversity and ecosystem service^1^. An assessment of the current status and trends in invasive alien species, their impacts and drivers, as well as their management and policy options is currently underway by the Intergovernmental Science-Policy Platform on Biodiversity and Ecosystem Services (IPBES) ^2,3^ The post-2020 target for invasive alien species by Parties to the Convention on Biological Diversity is under consideration^4^, and the 2030 Sustainable Development Goals include a target on the prevention of invasive alien species on land and in water systems^5^. These policy instruments require baseline data on the presence, trends of introduction and establishment, and impacts of invasive alien species.

Central to informing these processes, and to refining management and policy responses so that more rapid progress is made to achieving them, is information on the identity and distributions of the species concerned^6^. To support prioritisation of management efforts, information is also specifically needed on that subset of introduced species that negatively impact biodiversity and ecosystems^7^. Countries require such data, inter alia, to identify and keep track of novel invasions, prioritise investment in prevention and control efforts and meet multinational reporting obligations. Global aggregates of such data are essential (i) for countries to identify invasion risks from neighbouring regions, (ii) to identify gaps in taxonomic and distributional information, (iii) to prioritise and implement suitable prevention and surveillance measures, and (iv) for mandated policy reporting on this threat to biodiversity and ecosystems.

While substantive progress has been made over the recent decade to collate data on alien species and their distributions (e.g.^8–10^), these are largely taxon-specific, remain in the research arena and are not readily accessible to countries. Mostly they do not distinguish between alien species per se and the subset of these that harm biodiversity and ecosystems (i.e. invasive alien species, Table 1). In addition, no process exists for ensuring that policy-relevant information on the essential features of biological invasion (i.e. species identity, distribution and impact) is up to date and accessible, although numerous support tools are available^11^. Recent efforts ofthe GEO Biodiversity Observation Network (GEO BON) have highlighted the potential of Essential Biodiversity Variables (EBVs) for Species Populations to provide a foundation for the assessment and monitoring of the redistribution of species, including those considered invasive^12^. To progress work in this direction, a collaborative partnership process has been designed to support invasive alien species data updates^13^, and to support their EBV-based integration with other data sources to deliver indicators and other policy-relevant information^14^.

The purpose of GRIIS is to provide a range of products, including annotated checklists and checklist-based aggregations of information on introduced (Naturalised) and invasive alien species^1^ (Table 1, Fig. 1). GRIIS was introduced in 2018 using 20 exemplar Country Checklists^13^. The GRIIS Country Compendium presented here adds new value, inclusive of data of all countries that are Party to the Convention on Biological Diversity^15^ and the contiguous United States of America. The collation and publication of the Compendium presents a development advance over individual Country Checklists, including new data fields that enable cross-country and -taxon comparisons. It represents a significant advance from the estimated 52% of countries in 2010 with any form of listing of invasive alien species (not harmonised, necessarily accessible or taxonomically representative)^16^, to the GRIIS Country Compendium that now delivers comparable information across multiple taxa and for all countries. The Compendium is in a form that is findable, accessible, as interoperable as has been appropriate to achieve thus far, and reusable (FAIR^17^).

**Table 1.**
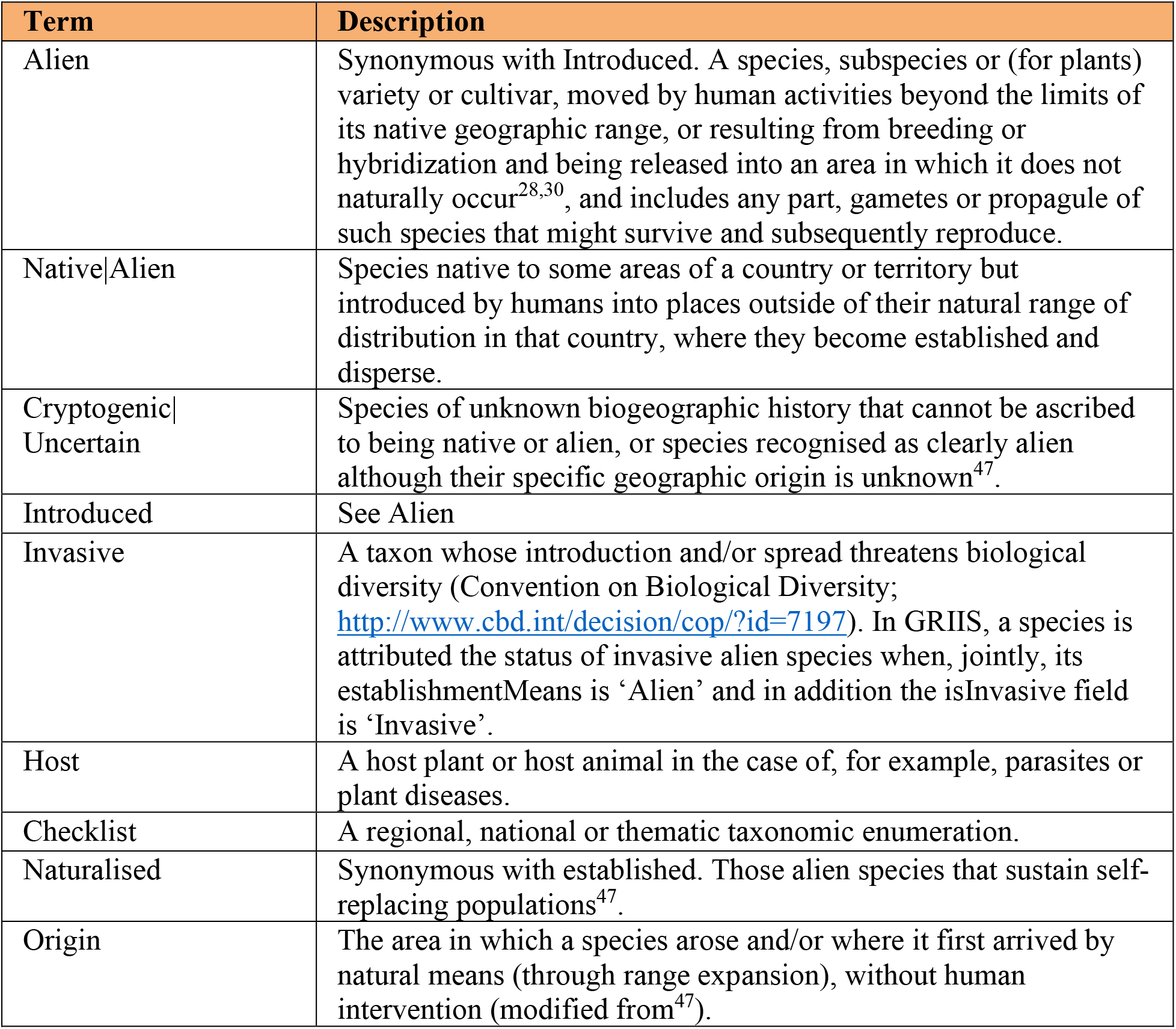
Glossary of general terms and their ecological meaning used in the Global Register of Introduced and Invasive Species (updated and modified from^13^)

**Figure 1.**
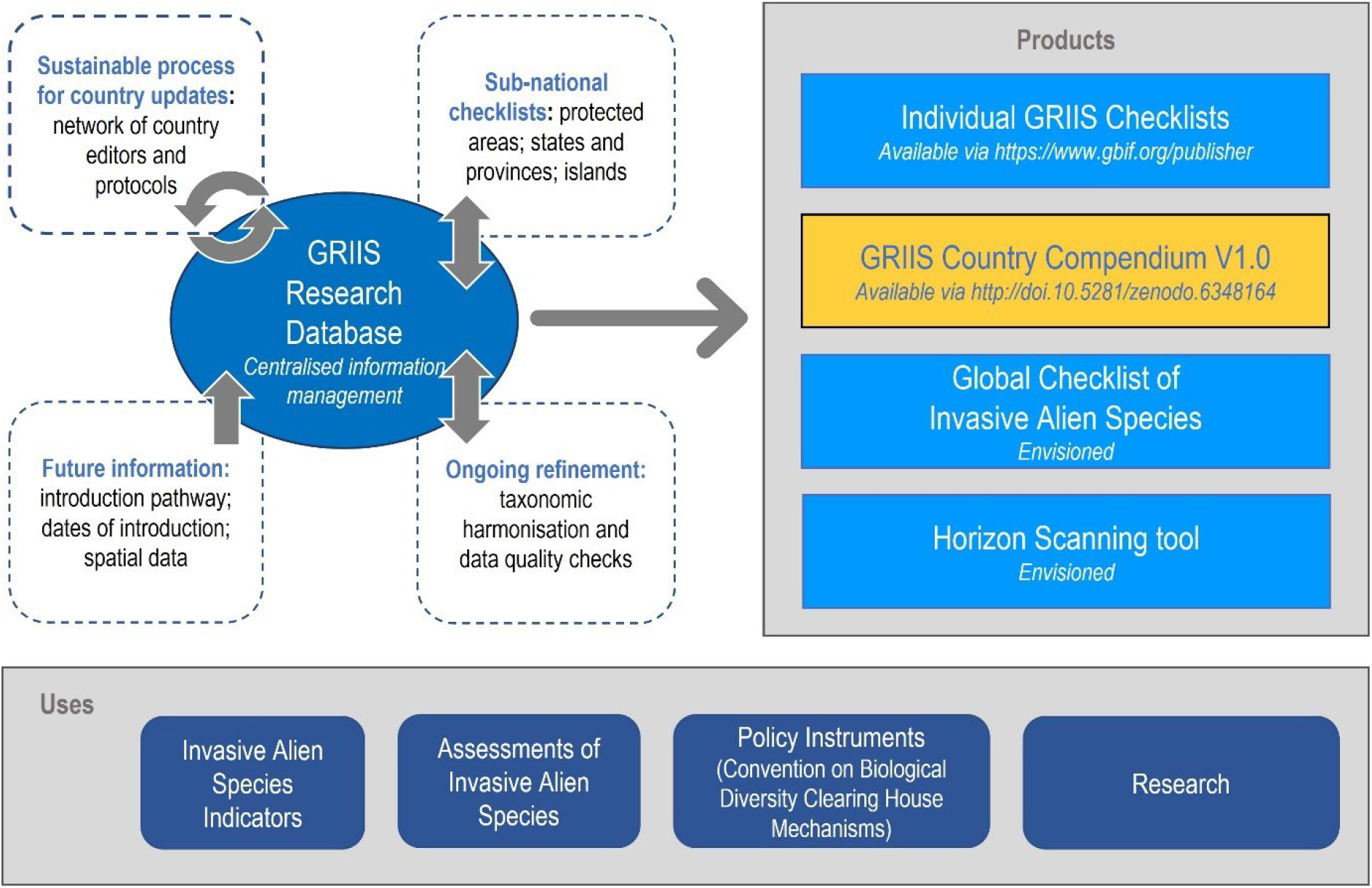
The GRIIS Country Compendium V1.0 (orange text box, right) shown as one of the key products of the Global Register of Introduced and Invasive Species (GRIIS) Research Database (centre left). Other data products are identified, including individual Checklists (including GRIIS Country Checklists^13^) available via GBIF.org, and the envisioned Global Checklist of Invasive Alien Species and Horizon Scanning tool. Grey arrows denote the envisaged interactive updates and information flows: Country updates are envisioned to occur over time, with the GRIIS Research Database supporting taxonomic harmonisation and data quality checks from information gathered through a global network of Country Editors. Each data product is then updated regularly as versioned snapshots of GRIIS. Uses of GRIIS (below) include use in global policy assessment and reporting and for research, such as by GEO BON, IPBES and the Convention on Biological Diversity Clearing House Mechanisms.

Here we publish and provide an overview of the GRIIS Country Compendium that provides a single, integrated dataset for research, assessment, monitoring and reporting at a global scale, including for example regional single and multi-species and taxon assessments and comparisons. It provides a key reference and baseline for future monitoring of the distribution and invasion status of the species^4^ involved, and for tracking progress toward global or local targets (Fig. 1).

## Methods

### GRIIS and the Country Compendium

The Global Register of Introduced and Invasive Species (GRIIS) arose following recognition of the need for a product of this nature in discussions on implementation of the Convention on Biological Diversity (CBD). In 2011, a joint work programme to strengthen information services on invasive alien species as a contribution towards Aichi Biodiversity Target 9 was developed (UNEP/CBD/SBSTTA/15/INF/14). The Global Invasive Alien Species Information Partnership (GIASI Partnership) was then established to assist Parties to the CBD, and others, to implement Article 8(h) and Target 9 of the Aichi Biodiversity Targets. The Conference of Parties (COP-11) welcomed the development of the GIASI Partnership and requested the Executive Secretary to facilitate its implementation (paragraph 22 of decision XI/28). In 2013, the development of GRIIS was identified as a key priority to be led by the IUCN ISSG and Partners built on a prototype initiated almost a decade earlier (Item 4, Report of the Global Invasive Alien Species Information Partnership, Steering Committee, 1st meeting Montreal, 15 October 2013)^18^.

GRIIS is a database of discrete checklists of alien species that are present in specified geographic units (including countries, islands, offshore territories, and protected areas) (Fig. 1). The GRIIS Country Compendium is a collation and key product that derives and is updatable from the working GRIIS Research Database that underpins this and other GRIIS products (Fig. 1). Individual checklists are published to GBIF through an installation of the Integrated Publishing Toolkit^19^ and hosted by the GBIF Secretariat (https://www.gbif.org/publisher/1cd669d0-80ea-11de-a9d0-f1765f95f18b). Exceptions include the Belgium (hosted by the Research Institute for Nature and Forest) and U.S.A checklists (hosted by the United States Geological Survey). Data are published as Darwin Core (dwc namespace) Archive files and the terms and structure follow that standard exchange format^20^.

The GRIIS Country Compendium is an aggregation of 196 GRIIS Country Checklists of which 82 % have been verified by Country Editors, along with revised and additional fields that enable global level analysis and country and taxon comparisons. Checklists for the 196 countries were combined into a single file (Table 3). A field was added to indicate which country the checklist belonged to, and the ISO3116-1 Alpha-2 and Alpha-3 country codes are included to facilitate dataset integration (see ‘Usage notes’) (Table 2). A field was also added to indicate the verification status of each checklist (Table 2). The ID field was renamed (originally ‘taxonID’ and now ‘recordID’), as the data now represent a country-level occurrence dataset containing multiple records per species, rather than checklist-type data that contains one record per species. In total, the data now include 18 fields as described in Table 2, encompassing taxonomic, location, habitat, occurrence, introduced and invasive alien status (see also Table 1). This publication represents a versioned, citable snapshot of the Compendium (Fig. 1) that is ready for analysis and integration with other data sources (e.g. workflow^21^ and ‘Example applications of the Compendium’ outlined further below).

**Table 2.** Fields and field terms in the GRIIS Country Compendium. An asterisk (*) denotes a field using a Darwin Core term^20^, while two asterisks indicate a term from the GBIF Species Profile extension^41^. The mapping of terms from the GRIIS Country Compendium to GRIIS Checklist Darwin Core Archive (available via GBIF) is shown in square brackets in the Field (‘taxonomicStatus’ has no equivalent in GRIIS).

| Field | Description | Terms in field |
| --- | --- | --- |
| recordID [taxonID] | GRIIS record identifier. |  |
| acceptedNameUsage* | The accepted name of the taxon, primarily according to GBIF and in some cases adjusted by GRIIS editors. |  |
| scientificName* | The name of the taxon as recorded during the data collection process, for example, the name by the country in question or in the original source material. |  |
| species | The accepted name of the taxon, given as a binomial name excluding subspecies and authorship information. The suffix "sp." is appended for genus-level records. |  |
| kingdom* | Higher level taxonomy, primarily according to GBIF and in some cases adjusted by GRIIS editors. Where not present or not relevant, “NOT ASSIGNED” is used. |  |
| phylum* |  |  |
| class* |  |  |
| order* |  |  |
| family* |  |  |
| taxonRank* | The rank of the taxon. | genus; species; subspecies; variety; form |
| isHybrid** | Whether the taxon is considered a hybrid. | true; false |
| country* | The name of the country as used by the United Nations. Note that country-level GRIIS checklists only cover the mainland of a country. |  |
| countryCode* | The alpha-2 code of the country according to the ISO 3166 standard. |  |
| countryCode_alpha3 | The alpha-3 code of the country according to the ISO 3166 standard. |  |
| habitat* | The dominant environment or environments occupied by a taxon. Multiple terms are delimited by a pipe character (\|). | terrestrial; marine; brackish; freshwater; Host |
| occurrenceStatus* | Presence or inferred absence of the taxon in the country. | present |
| establishmentMeans* | Introduced status of the taxon in the country. | Alien; Native\|Alien; Cryptogenic\|Uncertain |
| isInvasive** | A species is designated as ‘Invasive’ in a country using the systematic decision process outlined in Figure 2 of <sup>13</sup>. | Invasive; null |

**Table 3.**
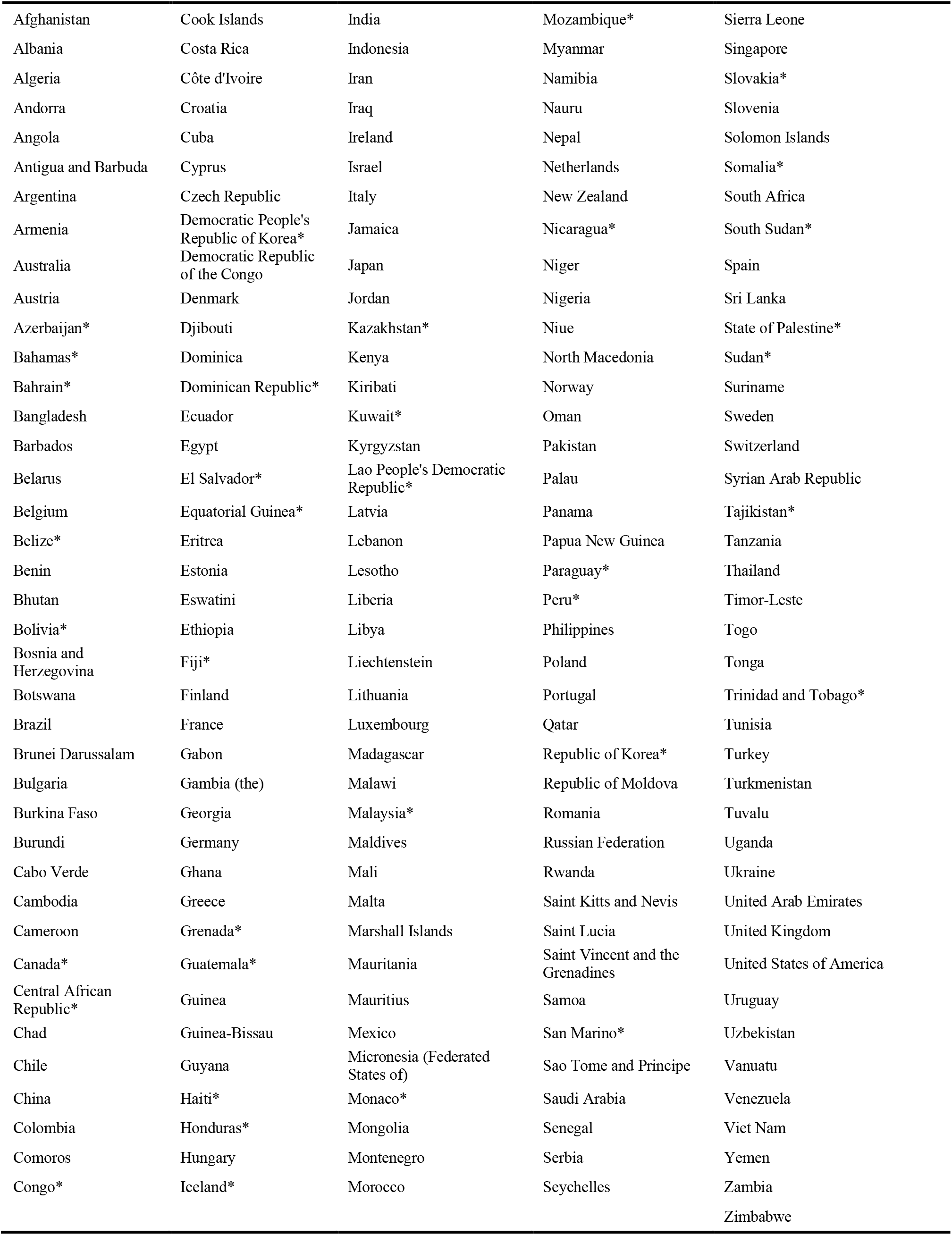
Countries in the GRIIS Country Compendium and their review status. Countries listed with an asterisk (*) have not (yet) been validated by country experts.

|  |  |  |  |  |
| --- | --- | --- | --- | --- |
| Afghanistan | Cook Islands | India | Mozambique* | Sierra Leone |
| Albania | Costa Rica | Indonesia | Myanmar | Singapore |
| Algeria | Côte d'Ivoire | Iran | Namibia | Slovakia* |
| Andorra | Croatia | Iraq | Nauru | Slovenia |
| Angola | Cuba | Ireland | Nepal | Solomon Islands |
| Antigua and Barbuda | Cyprus | Israel | Netherlands | Somalia* |
| Argentina | Czech Republic | Italy | New Zealand | South Africa |
| Armenia | Democratic People's Republic of Korea* | Jamaica | Nicaragua* | South Sudan* |
| Australia | Democratic Republic of the Congo | Japan | Niger | Spain |
| Austria | Denmark | Jordan | Nigeria | Sri Lanka |
| Azerbaijan* | Djibouti | Kazakhstan* | Niue | State of Palestine* |
| Bahamas* | Dominica | Kenya | North Macedonia | Sudan* |
| Bahrain* | Dominican Republic* | Kiribati | Norway | Suriname |
| Bangladesh | Ecuador | Kuwait* | Oman | Sweden |
| Barbados | Egypt | Kyrgyzstan | Pakistan | Switzerland |
| Belarus | El Salvador* | Lao People's Democratic Republic* | Palau | Syrian Arab Republic |
| Belgium | Equatorial Guinea* | Latvia | Panama | Tajikistan* |
| Belize* | Eritrea | Lebanon | Papua New Guinea | Tanzania |
| Benin | Estonia | Lesotho | Paraguay* | Thailand |
| Bhutan | Eswatini | Liberia | Peru* | Timor-Leste |
| Bolivia* | Ethiopia | Libya | Philippines | Togo |
| Bosnia and Herzegovina | Fiji* | Liechtenstein | Poland | Tonga |
| Botswana | Finland | Lithuania | Portugal | Trinidad and Tobago* |
| Brazil | France | Luxembourg | Qatar | Tunisia |
| Brunei Darussalam | Gabon | Madagascar | Republic of Korea* | Turkey |
| Bulgaria | Gambia (the) | Malawi | Republic of Moldova | Turkmenistan |
| Burkina Faso | Georgia | Malaysia* | Romania | Tuvalu |
| Burundi | Germany | Maldives | Russian Federation | Uganda |
| Cabo Verde | Ghana | Mali | Rwanda | Ukraine |
| Cambodia | Greece | Malta | Saint Kitts and Nevis | United Arab Emirates |
| Cameroon | Grenada* | Marshall Islands | Saint Lucia | United Kingdom |
| Canada* | Guatemala* | Mauritania | Saint Vincent and the Grenadines | United States of America |
| Central African Republic* | Guinea | Mauritius | Samoa | Uruguay |
| Chad | Guinea-Bissau | Mexico | San Marino* | Uzbekistan |
| Chile | Guyana | Micronesia (Federated States of) | Sao Tome and Principe | Vanuatu |
| China | Haiti* | Monaco* | Saudi Arabia | Venezuela |
| Colombia | Honduras* | Mongolia | Senegal | Viet Nam |
| Comoros | Hungary | Montenegro | Serbia | Yemen |
| Congo* | Iceland* | Morocco | Seychelles | Zambia |
|  |  |  |  | Zimbabwe |

### Population of data fields in GRIIS

The methods by which GRIIS is populated were described in 2018^13^ andare summarised in brief here. A systematic decision-making process is used for each geographic unit by species record to designate non-native origin and evidence of impact (see Fig. 2 in Pagad et al.^13^). Comprehensive searches are undertaken for each country. Records are included from the earliest documented to the most recent accessed record prior to the date of the latest published checklist version. Information sources include peer-reviewed scientific publications, national checklists and databases, reports containing results of surveys of alien and invasive alien species, general reports (including unpublished government reports), and datasets held by researchers and practitioners^13^. A log of the changes to each checklist^22^ is available on the GBIF IPT^23^, with the changes to the Belgium checklist available at the INBO IPT^24^. The most up to date version of each checklist is thus available via GBIF.org, as is a list of all GRIIS checklists at GBIF.org^23^.

**Figure 2.**
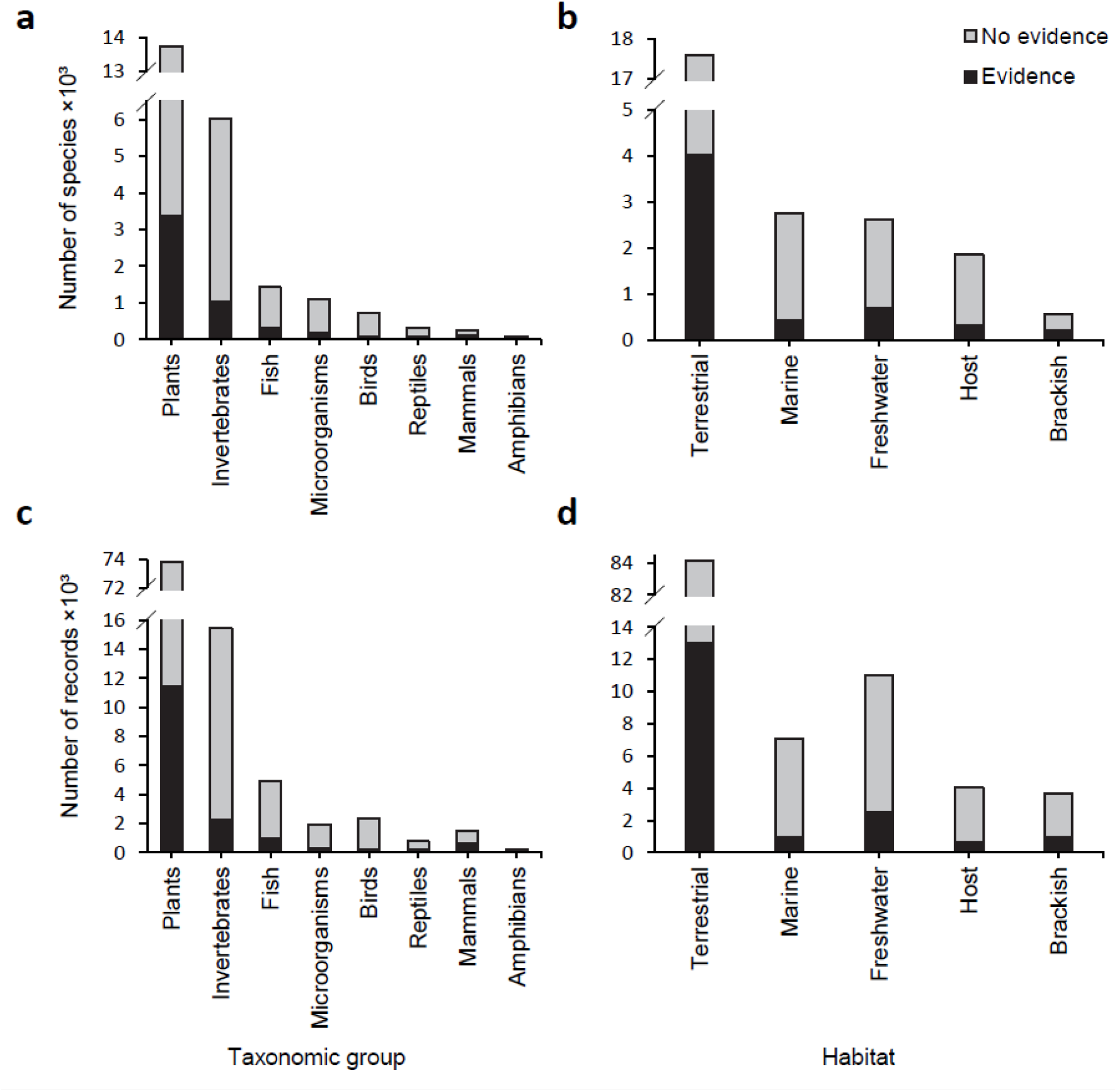
Summary of data in the GRIIS Country Compendium. Number of Introduced and Invasive species by major taxonomic group (a) and habitat (b). Number of records per major taxonomic groups (c) and habitat (d). The number of species and records associated with invasion impact (i.e. isInvasive) are shown in black. Note different y-axis scales in each case.

Introduced species of all taxonomic groups are considered for inclusion in GRIIS. Environments include terrestrial, freshwater, brackish, marine and also host (i.e. for species that not free-living) (Table 1). Typically, GRIIS records are at the species level, but in some cases, other ranks are more appropriate including infraspecies (including forms, varieties and subspecies). A separate field is provided for hybrids (Table 2). Where species are present and both native to parts of a country and alien in other parts of the country, their dwc:establishmentMeans is included as Native|Alien (Tables 1,3)^25^. If there is limited knowledge about the Origin of the species, its dwc:establishmentMeans is included as Cryptogenic|Uncertain (Tables 1, 3). The habitat information in GRIIS (Table 2) is sourced from taxon and region-specific databases such as WoRMS (World Register of Marine Species), FishBase, Pacific Island Ecosystems at Risk, and the USDA Plants Database.

Two types of evidence are considered to assign a species by country record as invasive (Table 1) (sensu^13^) (Table 1): (i) when any authoritative source (e.g. from the primary literature or unpublished reports from country/species experts), describe an environmental impact, and/or (ii) when any source determines the species to be widespread, spreading rapidly or present in high abundance (based on the assumption that cover, abundance, high rates of population growth or spread are positively correlated with impact)^26,27^. Each record is assigned either invasive or null in the isInvasive field to reflect the presence of evidence of impact, or absence of evidence of impact (note not ‘evidence of absence of impact’), for that species by country record (Table 2). In the future this information may be supplemented with impact scores^28–30^.

Finally, a draft checklist is sent to Country Editors for validation and revision (see Technical Validation).

### Taxonomic harmonization and normalization

The use of different synonyms across countries to refer to the same taxonomic concept is frequent^31^. The species in each country checklist were thus harmonised against the GBIF Backbone Taxonomy^32^. The names in each checklist were matched using a custom script that integrates with the GBIF API^33^, and the accepted name, taxon rank, status and higher taxonomy (Table 2) were also obtained at this stage. Spelling and other errors in assigning species authorship were corrected where appropriate.

To validate the taxonomic harmonisation, every name variant present in the GRIIS Country Compendium was checked against the GBIF Backbone Taxonomy using the API^11^. A unique list of names from both the reported name and the accepted scientific name (Table 2) was checked (n=32 443). Over 95% of names across all kingdoms matched exactly at 98% or greater confidence (Table 4). All names that were below 98% confidence or had a match type other than ‘Exact’ were checked and modified if appropriate to do so. Of the non-matches (n=253, those with a match type of ‘None’), most were formulaic hybrid names of plants and animals (~62%), which are not officially supported by GBIF^34^. The remaining non-matches were names of mostly plants (17%), but also animals (8%), viruses (8%) and chromists (3%).

**Table 4.** Taxonomic matching results (percentages) by Kingdom using the GBIF Backbone Taxonomy (GBIF Secretariat 2021). Results are shown as a percentage of the total number of unique names in the GRIIS Country Compendium (such that the table total sums to 100% of all unique names in GRIIS), including names from both scientificName and acceptedNameUsage fields. Results are split by the Kingdom of the matched species. Protozoa excluded (0.03% of names matched exactly at 98% or greater confidence).

| Match type | Animalia | Bacteria | Chromista | Fungi | Plantae | Total |
| --- | --- | --- | --- | --- | --- | --- |
| Exact – 98% or greater confidence | 31.32 | 0.13 | 1.69 | 1.89 | 59.93 | 95.00 |
| Exact – below 98% confidence | 0.34 | 0.00 | 0.03 | 0.02 | 1.85 | 2.23 |
| Fuzzy – partial match | 0.17 | 0.00 | 0.00 | 0.01 | 0.20 | 0.38 |
| Higher rank – name matched to a taxon at a level higher than species | 0.48 | 0.01 | 0.16 | 0.00 | 0.96 | 1.61 |
| None – no match | *NA* | *NA* | *NA* | *NA* | *NA* | 0.78 |

### Data summary

There are currently ~23 700 species represented by 101 000 taxon-country combination records, across 196 countries in the GRIIS Country Compendium. All raw numbers are provided to the nearest order of magnitude to reflect the taxonomic uncertainty and dynamic nature of GRIIS (see ‘Known data gaps and uncertainties’). The vast majority of records are at the species level (97.6%), with the remaining present as subspecies (1.7%), varieties (0.6%), genera (0.1%) and forms (<0.1%). In addition, approximately 0.5% of records are either named or hybrid formulae. For the purpose of providing an overview of the content of the Compendium, we counted species using the accepted species name in the ‘species’ field (for this purpose not differentiating by taxonRank) (Table 2).

Most country checklists include between 100-1000 species (137); 36 checklists have < 100 species and 23 checklists with > 1000 species (minimum = 5; maximum values = 5649). On average the number of species per country is 516 (721 s.d.), with the fewest records for San Marino, South Sudan and Monaco (<12) and most for the U.S.A, France and Australia (>2900).

The taxonomic and habitat coverage in the Compendium is broad, with 55 phyla, 149 classes, 630 orders and 2305 families represented across terrestrial, marine, freshwater and brackish habitats (Fig. 2). Those currently represented include Animalia, Bacteria, Chromista, Fungi, Plantae, Protista (Protozoa), and Viruses. Over 98% of records have establishmentMeans as simply alien, whereas 1% of records are Cryptogenic|Uncertain and 1% Native|Alien (Tables 1, 3).

As expected for invasive alien species, the Compendium is dominated by plant and invertebrate species and records (Fig. 2a, c). Overall, 16% (n=16 221) of all records are associated with impact (isInvasive, Table 2), and 22% of species (n=5199) are associated with evidence of impact in at least one country. Terrestrial habitats have most species and records associated with evidence of impact, followed by freshwater and marine species and records (Fig. 2b, d). Approximately 15% of plant (n=11,436) and 16% of invertebrate (n=2300) species by country records are associated with evidence of impact, whereas mammal (42%, n=644) and amphibian (34%, n=84) records have higher percentages of evidence of impact at the country level. Over half of the species in the Compendium are recorded from only a single country, with 10.5% present in more than 10 countries (Fig. 3a). As expected, the numbers of ‘isInvasive’ records per species are lower than the total number of records per species, although species with many country occurrence records do tend to have many ‘isInvasive’ status annotations (Fig. 3b).

**Figure 3.**
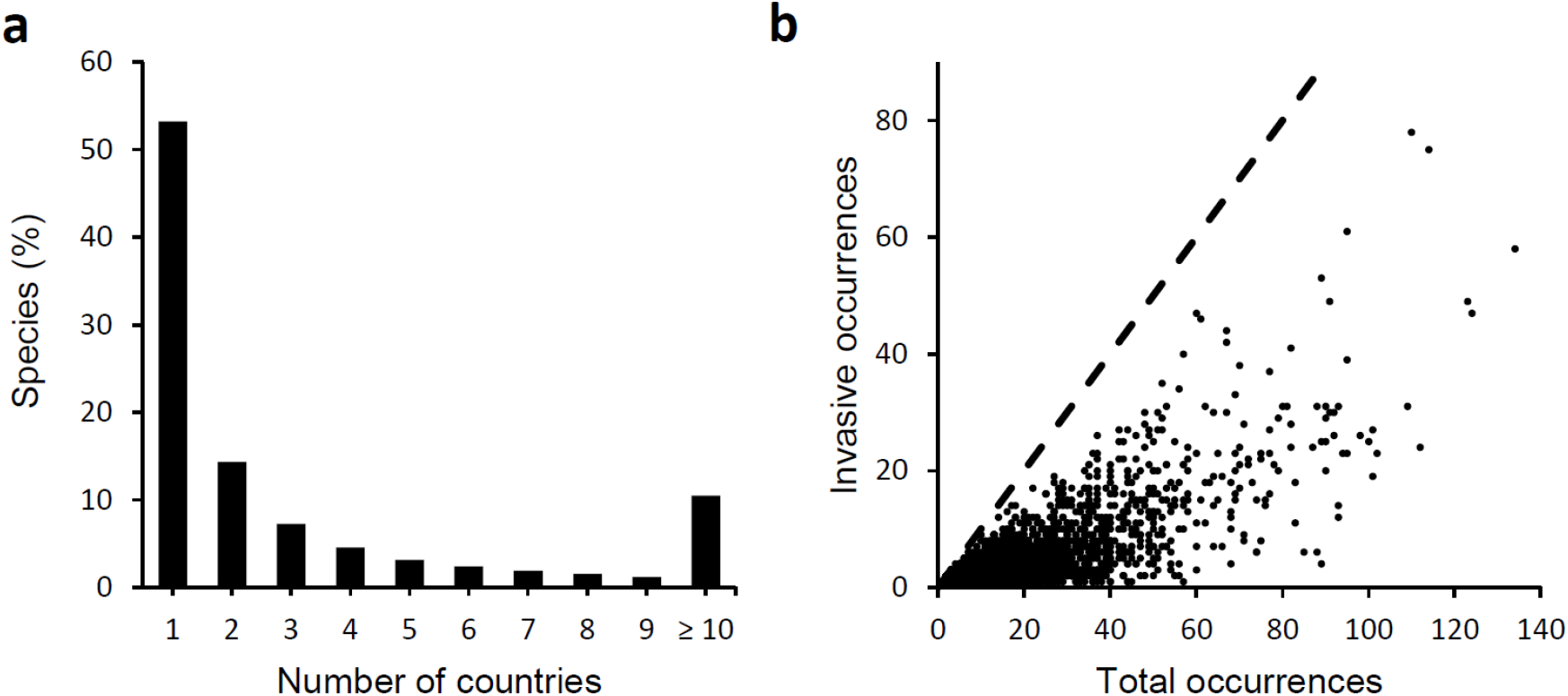
(a) Frequency of all species (n= ~ 23,600) across countries (n=196), showing that the majority of species are reported for only a single country. (b) The relationship between the total number of occurrences (i.e. taxon-country records) and number of occurrences associated with evidence of impact for each species.

### Example applications of the Compendium

The GRIIS Country Compendium enables the first multitaxon assessment of the most widely distributed subset of introduced species that harm biodiversity and ecosystems. These ‘worst’ invasive alien species (in terms of realised impact across most countries), including wild sage, black rat, common carp, red-eared slider turtle, American bullfrog, common myna, sea walnut and giant African snail(Table 5). To demonstrate key envisaged uses of the Compendium, it has been incorporated in Map of Life enabling: (1) multitaxon mapping of richness patterns in invasive alien species (e.g. Fig. 4), and (2) the combination, at the species level, of alien information along with point observations, expert defined ranges and other types of data describing species distributions worldwide (Fig. 5). This facilitates model-based integration of the Compendium data with environmental data to assess changes and trends in species distributions. The Compendium also enables the consideration of gaps and inferred absences of species at the country scale useful for the purpose of updating country checklists, informing risk assessments and for research on the drivers of invasive alien species spread and establishment^21,35^. For example, occurrence records for the red eared slider turtle in East Africa, the Middle East and Australia (Fig. 5) suggest that GRIIS Country Checklists for the countries concerned may need to be updated, triggering an update to the Compendium. A growing number of publications now cite use of multiple GRIIS checklists, accessed through GBIF, by use of Digital Object Identifiers^36^. The Compendium will now streamline and facilitate such multi-checklist applications.

**Table 5.** Top five species per taxonomic group by number of country by species records (occurrences) flagged as invasive in the GRIIS Country Compendium. Invertebrates split by ‘habitat’ field; with parasitic (host) and brackish invertebratesomitted. Invertebrates in multiple environments were assigned marine, terrestrial, or freshwater in that order. Common name sourced from NCBI (https://www.ncbi.nlm.nih.gov/) then GISD (http://www.iucngisd.org/gisd/ISSG_n.d.)

| Group | Species | Common name | Invasive occurrences | Total occurrences |
| --- | --- | --- | --- | --- |
| **Plants** | *Lantana camara* | Wild sage | 78 | 110 |
|  | *Eichhornia crassipes* | Water hyacinth | 75 | 114 |
|  | *Leucaena leucocephala* | White popinac | 58 | 134 |
|  | *Chromolaena odorata* | Bitter bush | 47 | 60 |
|  | *Ricinus communis* | Castor bean | 47 | 124 |
| **Mammals** | *Rattus rattus* | Black rat | 61 | 95 |
|  | *Mus musculus* | House mouse | 53 | 89 |
|  | *Rattus norvegicus* | Norway rat | 49 | 91 |
|  | *Felis catus* | Domestic cat | 40 | 57 |
|  | *Sus scrofa* | Pig | 34 | 56 |
| **Fish** | *Cyprinus carpio* | Common carp | 49 | 123 |
|  | *Gambusia holbrooki* | Eastern mosquitofish | 42 | 67 |
|  | *Oreochromis niloticus* | Nile tilapia | 30 | 92 |
|  | *Oreochromis mossambicus* | Mozambique tilapia | 25 | 89 |
|  | *Oncorhynchus mykiss* | Rainbow trout | 25 | 100 |
| **Reptiles** | *Trachemys scripta* | Red-eared slider turtle | 31 | 81 |
|  | *Hemidactylus frenatus* | Common house gecko | 13 | 32 |
|  | *Hemidactylus mabouia* | Tropical house gecko | 8 | 26 |
|  | *Iguana iguana* | Common green iguana | 5 | 18 |
|  | *Chelydra serpentina* | Common snapping turtle | 4 | 12 |
| **Amphibians** | *Lithobates catesbeianus* | American bullfrog | 25 | 42 |
|  | *Rhinella marina* | Marine toad | 14 | 30 |
|  | *Xenopus laevis* | African clawed frog | 10 | 16 |
|  | *Triturus carnifex* | Italian crested newt | 4 | 6 |
|  | *Eleutherodactylus johnstonei* | Johnstone's robber frog | 3 | 12 |
| **Birds** | *Acridotheres tristis* | Common myna | 22 | 46 |
|  | *Columba livia* | Rock pigeon | 20 | 90 |
|  | *Corvus splendens* | House crow | 17 | 34 |
|  | *Passer domesticus* | House sparrow | 14 | 55 |
|  | *Psittacula krameri* | Rose-ringed parakeet | 13 | 50 |
| **Marine invertebrates** | *Mnemiopsis leidyi* | Sea walnut | 18 | 29 |
|  | *Magallana gigas* | Pacific oyster | 15 | 52 |
|  | *Ficopomatus enigmaticus* | Australian tubeworm | 15 | 31 |
|  | *Acanthaster planci* | Crown-of-thorns starfish | 14 | 18 |
|  | *Eriocheir sinensis* | Chinese mitten crab | 14 | 26 |
| **Terrestrial invertebrates** | *Lissachatina fulica* | Giant African snail | 31 | 53 |
|  | *Tapinoma melanocephalum* | Ghost ant | 28 | 71 |
|  | *Pheidole megacephala* | Big-headed ant | 27 | 52 |
|  | *Aedes albopictus* | Asian tiger mosquito | 24 | 70 |
|  | *Solenopsis geminata* | Tropical fire ant | 19 | 35 |
| **Freshwater invertebrates** | *Corbicula fluminea* | Asian clam | 23 | 37 |
|  | *Dreissena polymorpha* | Eurasian zebra mussel | 21 | 35 |
|  | *Procambarus clarkii* | Red swamp crayfish | 19 | 40 |
|  | *Pacifastacus leniusculus* | Signal crayfish | 19 | 27 |
|  | *Potamopyrgus antipodarum* | New Zealand mudsnail | 15 | 37 |
| **Microorganisms (including macroalga)** | *Vibrio cholerae* | Asiatic cholera | 17 | 22 |
|  | *Aphanomyces astaci* | Crayfish plague | 13 | 16 |
|  | *Batrachochytrium dendrobatidis* | Amphibian chytrid fungus | 10 | 25 |
|  | *Undaria pinnatifida* | Asian kelp | 10 | 17 |
|  | *Ophiostoma novo-ulmi* | Dutch elm disease | 10 | 17 |

**Figure 4.**
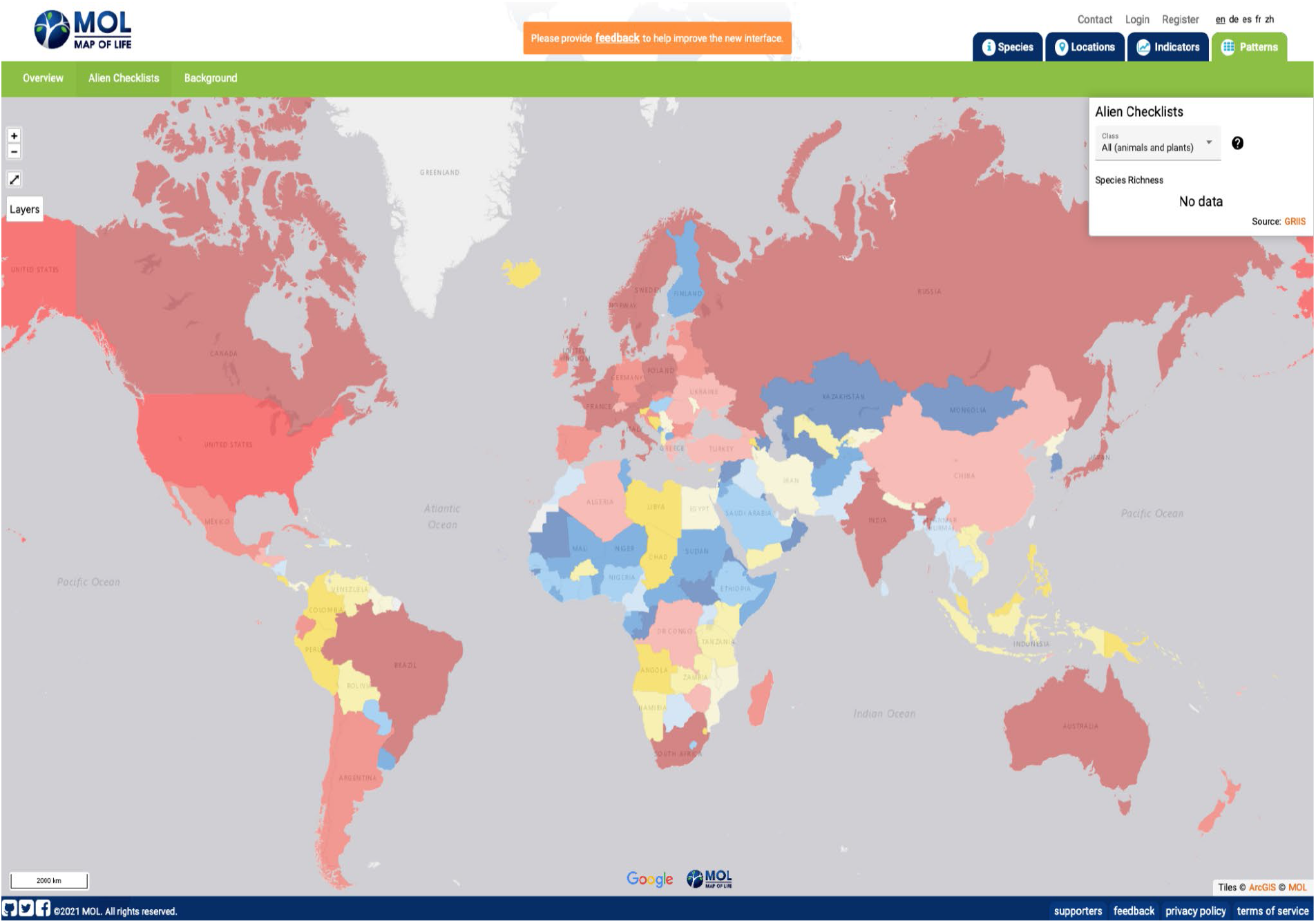
Use of GRIIS Country Compendium data to map the richness of Introduced and Invasive species, here for vertebrate and plant species across countries (n=196) and taxonomic groups (~5600 species). The aggregated data here include all records from amphibians, reptiles, birds, mammals and plants that were identified at the species level and exclude infraspecies. Countries or regions in grey not represented in GRIIS. Map compiled in Map of Life and visible at: https://mol.org/patterns/aliens

**Figure 5.**
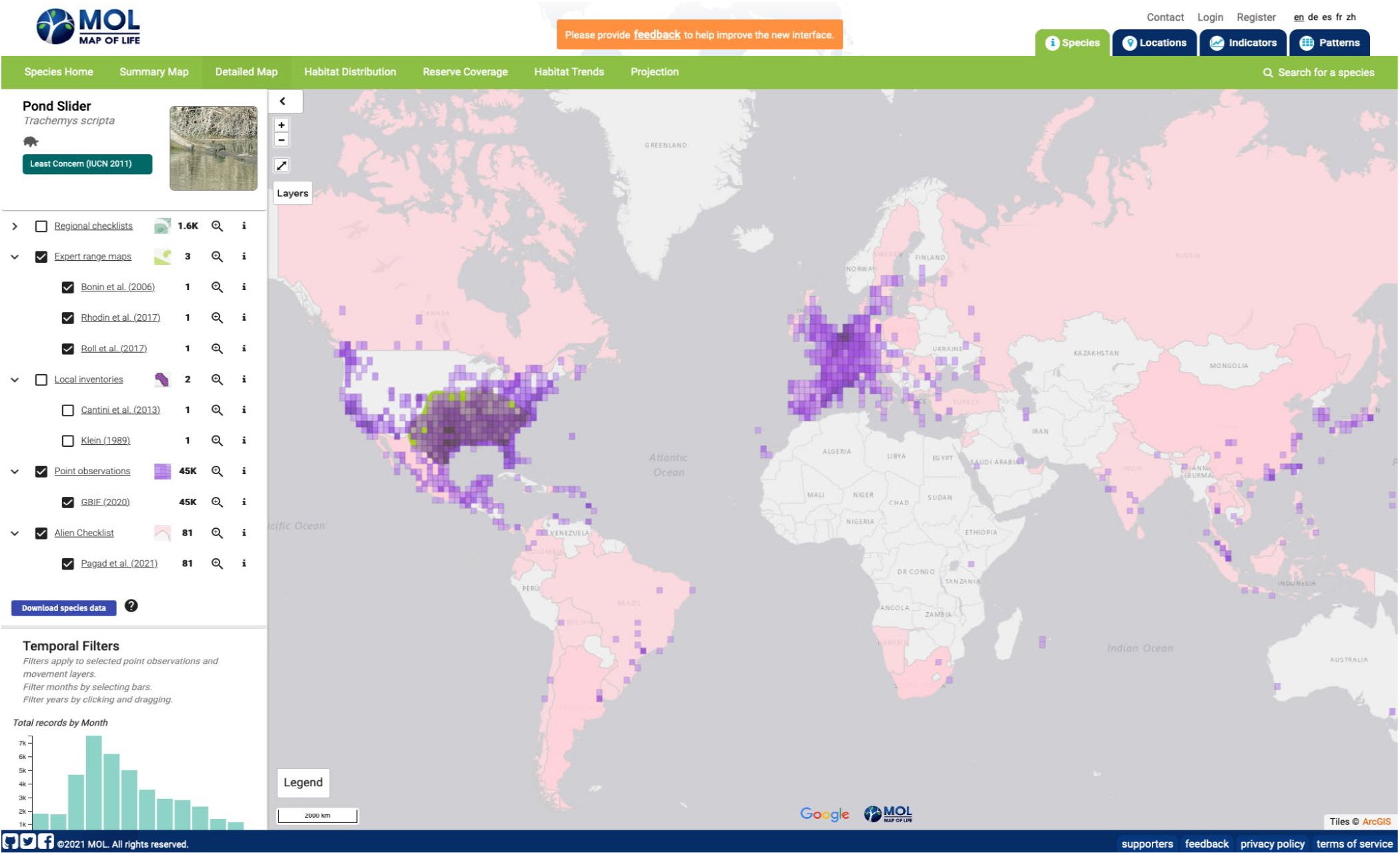
Example of a species page illustrating the complementary information value of GRIIS data, species occurrence records (mediated through GBIF) and native range information (in green). The example here is of the invasive alien Red eared slider turtle (*Trachemys scripta*, n=65), which, based on the Compendium data, has the most widely distributed evidence of impact outside of its native geographic range of any amphibian species (Table 5). GRIIS taxon-country records (in pink) are overlaid with individual occurrence records (purple) and include expert native range information (green). Map compiled in Map of Life and visible at: https://mol.org/species/map/Trachemys_scripta

The Compendium will further facilitate international research avoiding the need for repeated multi-country data integration by individual researchers. In support of surveillance and information currency, it provides a mechanism for generating automated alerts of risks of new incursions by analysing presence in neighbouring countries, especially through development of further workflows reporting new occurrence data published through GBIF. The Compendium may be used in decision support, contributing to invasion pathway and impact analyses and horizon-scanning exercises. In addition to the GRIIS Country Checklists available to national governments through the CBD Clearing House Mechanism (CHM) and national CHMs (www.cbd.int/countries), the GRIIS Country Compendium provides a foundation for long-term reporting on progress towards meeting invasive alien species-relevant goals and targets under the Convention on Biological Diversity, the Sustainable Development Goals and regional or thematic goals and targets. For example, the habitat data included in the Compendium could be disaggregated and used by relevant conventions, such as The Ramsar Convention on Wetlands and the Convention on Migratory Species. The Country Compendium data were also used in the development of the Global Biodiversity Outlook GBO5 and are being used in the IPBES Invasive Alien Species Assessment currently underway.

The Compendium provides the baseline used in the development of the invasive alien species headline indicator that measures trends in introduction events. It provides the key information source for Essential Biodiversity Variable-based indicators of invasion, including rate of spread, impact and status information^35^. This includes an indicator for reporting against the draft post-2020 Global Biodiversity Framework (CBD/WG2020/3/3) Target 5 and associated Headline Indicator – Rate of Invasive Alien Species Spread (CBD/WG2020/3/3/ADD1), enabling the calculation of baseline values at different scales, including the country level. Aggregated data from GRIIS, now published as the GRIIS Country Compendium, have thus already supported multiple research and key policy activities.

### Known data gaps and uncertainties

One of the principal objectives of GRIIS is to provide a geographically and taxonomically comparable source of information on invasive alien species. The intent is to collate the taxonomy and use the same approach to designating every taxon as alien, Native Alien, Cryptogenic, invasive or not (Table 1). These factors have previously been the cause of major uncertainty in collations of information on alien species. Nonetheless, a number of known gaps and uncertainties remain in the Compendium.

### Taxonomy

Sources of taxonomic uncertainty include multiple accepted names (acceptedNameUsage) for the same species concept - including author variations, lexical variants and potentially different taxonomic interpretation from the source. For example, the bay barnacle is associated with both *Balanus improvisus* and *Amphibalanus improvisus*. There are also multiple scientificName variants for individual acceptedNameUsages. No attempt has been made to edit or match species authorities or reference literature defining the taxon concepts. The species-level prevalence of lexical variants, synonyms and infraspecific names used across countries in the scientificName field is 20%, i.e. ~ 4740 species with more than one scientific name used, ~1520 species with more than 2 names, 14 species with more than 10 names and a maximum of fifteen scientificName variants for a single species. Overall, 4.5 % of unique species names are associated with an infraspecific name. The average percentage of such cases across species groups is ~14%, with the lowest percentage for birds (~9 %) and the highest for plants (~ 27 %) (microorganisms not considered).

The taxonomic information in the Compendium thus includes sources of uncertainty in common with all interspecific inventories of taxonomic information^37^, including different usage of synonyms across countries that could result in the inclusion of the same species as multiple taxa^38^. A shortage of editors specializing in particular taxonomic groups within countries is a recurring problem, as is the time necessary for editors to conduct and return reviews of draft checklists. Nonetheless, the most appropriate approach for harmonising taxonomy is determined by the intended use of the dataset. The Compendium provides both harmonised and un-harmonised taxonomic data in different fields to enable users to select the appropriate data for their use case. The accepted name usages provided are not necessarily the most taxonomically up to date usage, or necessarily the preferred usage of a particular country, but this field is included to enable comparisons across countries. For instance, Map of Life used harmonized taxonomic data excluding all infraspecies level records (i.e., subspecies, hybrids, varieties) to integrate GRIIS data with other types of data describing the distribution of species (e.g., species occurrence data mediated through GBIF, or expert range maps) (Fig.5)

### Completeness

Invasive alien species checklists are inherently dynamic, and indeed the value they embody is in the changes in the inclusion and exclusion of species over time. Alien and invasive alien species databases and lists are by nature dynamic for multiple reasons, including the population and range dynamics of alien species, the time lag that often occurs between establishment and spread of alien species outside of their native ranges, and time lags between such events and the quantification and documentation of their impacts^39^. Other biases include those well known for alien species, including geographic bias and taxonomic gaps, language barriers to information access and collation, and inconsistent use of invasion terminology across information sources^39,40^. In addition, potential sources of uncertainty and error include (i) misidentification of particular species in particular countries, (ii) species listed as being present that have subsequently become extinct or been eradicated, (iii) species listed as present based on in-country occurrence records but where the species has not established, and (iv) cases where particular taxonomic groups are undergoing rapid or regular taxonomic revision.

GRIIS aims to be a list, as complete as possible, of those alien species both confirmed to be present and also associated with evidence of impact in one or more countries, i.e. of ‘true presences’ of introduced species with evidence of impact. It is less complete as a list of true presences of alien species for which no evidence of negative impacts on biodiversity and ecosystems exists. In other words its focus is on the subset of alien species most likley to be a policyand management priority^7^. GRIIS should be regarded and appreciated as an evidence-based information repository designed for the purpose of improving and monitoring change in alien and invasive status and properties. Acknowledging the uncertainties and the dynamic properties of these data (in some cases inherent and in other cases by design), by providing a transparent, traceable process of data collation, the GRIIS Compendium provides a platform for delivering, refining and updating information on invasive alien species needed for global reporting, a baseline for ongoing updates, and a platform for sustainable delivery of this information (Fig. 1).

### Data Records

The static, versioned GRIIS Country Compendium (V1.0) is available as a ZIP archive, containing a UTF-8 Comma Separated Values file and associated metadata file [https://doi.org/10.5281/zenodo.6348164, 19 April 2022, 19 April 2022]. At the same time, dynamic individual country checklists are available, as outlined^13^, through GRIIS.org. Each row in the Compendium represents the presence of a taxon in a country. Where possible, we present the original name usages in ‘ scientificName (the name used by the original source or by the country itself), to allow users to harmonise the names according to their need (Table 2). We also provide ‘acceptedNameUsage, which is the accepted name recorded during the data aggregation process (Table 2). Finally, we provide ‘species’, which is the accepted binomial name (ie. excluding infraspecies epithet) which is useful for comparing species across countries where differentiating infraspecies is not desired (Table 2). While field names are consistent with Darwin Core terms^20,25^ and appropriate extensions^41^ as far as practicably possible, the data also include additional non-standard terms for accuracy, clarity, and ease-of-use (Table 2). GRIIS taxon-country records are publicly available for visualisation in Map of Life species page under the ‘Detailed Map’ tab (e.g., https://mol.org/species/map/Trachemys_scripta, Fig. 5) for any given species included in the Compendium. Aggregated records by species and country are also available for visualization in the patterns page (https://mol.org/patterns/aliens/).

### Technical Validation

The validation of species present in a country and evidence of their impact in that country followed the same process outlined in Pagad et al.^13^. Validation involves the harmonization of taxonomy, assigning the habitat for each species, checking that the interpretation of the taxonomic status of the species, its Origin, occurrence status and evidence of impact (Tables 1 and 2) are correctly recorded. In total, 82% of checklists in the GRIIS Country Compendium were reviewed prior to publication (review has not yet proved possible for the remaining 18% of countries) (Table 3). Country Editors reviewed the accuracy and completeness of records based on their specific expertise (Table 3), and the information and knowledge available to them (see^13^ for the details of this process). These Country Editors are authors of the published Country Checklists.

### Usage Notes

#### End-user harmonisation

If the end-user intends to harmonise the names provided in GRIIS, using the ‘scientificName’ field is recommended. The names provided in the ‘species’ and ‘acceptedNameUsage’ fields have already undergone some harmonisation to the GBIF Backbone Taxonomy.

#### Mapping GRIIS and geographic boundaries

If the end-user intends to map the data, or use the ISO 3166 country codes to map to other data sources, they should be aware that the country-level GRIIS Checklists provided here generally pertain to the mainland of the country only with some exceptions, e.g., the United States that includes the contiguous 48 states. Spatial data on geopolitical boundaries is increasingly available to support mapping of GRIIS data at sub country level differentiating mainland and offshore territories^42^.

#### Mapping the Country Compendium to GBIF Darwin Core Archives

The GRIIS Country Compendium includes four fields not present in Darwin Core Archives of GRIIS Checklists (Table 2). The ‘taxonID’ field was renamed ‘recordID’ to clarify that it is a unique identifier for the GRIIS record, rather than an identifier for the taxonomic concept. The ‘countryCode_alpha3’ field was added to further enable dataset integration. The ‘species’ field was added to enable data analysis, while the ‘isHybrid’ field was added to provide additional context to the record.

#### Maintenance, updates and expansion

GRIIS is being continually improved and updated, triggered by updates from Country Editors (see Technical Validation), new scientific literature, feedback from users, and/or analysis of occurrence data published through GBIF.org as a source of evidence that an alien species has not been currently included (Fig. 1). Any updates follow a process of validation and verification. It is envisioned that future updates will be released, with additional species and corrections (Fig. 1).

Ongoing, versioned updates to the GRIIS Country Compendium V1.0 can occur as a result of (i) improved knowledge of alien species and their impacts in countries, and (ii) the dynamics of invasive alien species distributions as a consequence of ongoing range expansions, as well as reductions as a consequence of control efforts. Subject to the availability of funding, engagement with Country Editors and incremental updates are planned on an on-going basis (e.g., data available as a result of new research, surveys, publications, changes in the status of species, new incursions, successful and proved eradications, and occurrence data published from multiple sources through GBIF). Scheduled major updates are planned in such a way that a review and major update of every GRIIS Country Checklist is completed once every two years, with automated integration into the Compendium (Fig. 1).

The intention is to continue to add key data fields to the published Compendium as they are completed or verified. Envisaged GRIIS developments to follow include: publication of known dates of introduction or first report for species; coded source information for each record; checklists for all European Overseas territories; known pathways of country by species records; impact mechanisms for species; mechanisms of impact aligned with EICAT^28–30^; other geographic units encompassed by GRIIS (islands, offshore territories, and protected areas); spatial data designating the relevant boundaries (for example combining national and sub national information from the Database of Global Administrative Areas (gadm.org) with coastline and island information from remote-sensed, high-resolution global shoreline vector (GSV-USGS^42^). Following a process of taxonomic harmonisation, a Global Checklist of Invasive Alien Species will be published as a further product of GRIIS, flagging individual alien species known to be invasive anywhere in the world (Fig. 1).

#### GRIIS Citation guidelines

A number of data citation options are available to cover reference to GRIIS work and data: (i) To refer to the Compendium and latest progress by GRIIS, cite this paper; (ii) to cite the GRIIS.org portal, cite^13^; (iii) to cite national or subnational datasets as available through GBIF.org, refer to the Citation sections at the bottom of each checklist dataset page, e.g. for Australia cite^43^, and see GBIF citation guidelines^44^; (iv) to cite GRIIS based data products from Map of Life, cite^45^; (v) to cite GRIIS checklists as national reference points for the Convention of Biological Diversity, cite^13^. It is also possible to query combinations of individual records and cite the resulting data downloads, access data via GBIF API^33^, or cite the DOI for derived datasets^46^.

#### Ongoing support for the development and maintenance of GRIIS

The daily maintenance and management of GRIIS is based on a low-cost model and currently includes a team of two, with expert input from key ISSG members. The IUCN ISSG researcher and practitioner networks that have been built over two decades are key supporters and contributors. The initial development of the country checklists as well as their integration with the GBIF data infrastructure have been supported through time-limited grants provided by, among others, the CBD and GBIF. However, the current model based on project funding is proving to be an impediment to provision of data and information on real time to support informed decision making. Maintenance and regular updating of GRIIS, keeping abreast of evolving data standards, with sufficient funding and human resources, are clearly essential if its utility as an information source is to be continued into the long term. The GRIIS team is working with relevant partners to solicit support from governments and institutions to enable this critical work to continue in a sustainable way.

## Supporting information

Metadata for GRIIS Country Compendium dataset

## Acknowledgements

GRIIS was developed and managed by the IUCN SSC Invasive Species Specialist Group (ISSG), with co-funding from the European Union through the Secretariat of the Convention on Biological Diversity within the framework of the Global Invasive Alien Species Information Partnership (GIASIPartnership). We are exceptionally grateful to the over 400 number of Country Editors who contributed to verifying draft Country Checklists and particular taxonomic groups (Country Editor information is available and associated with GRIIS Country Checklists at: https://www.gbif.org/publisher/cdef28b1-db4e-4c58-aa71-3c5238c2d0b5). Major support from the Global Biodiversity Information Partnership (GBIF) enabled the completion of the global coverage and publication of GRIIS country checklists. We thank Joe Miller, Marcus Döring, Daniel Noesgaard, Tim Robertson, Andrea Hahn, and the Data Products team of GBIF for comments on the manuscript and support with the repository. MM acknowledges GEO BON support of this initiative, and Australian Research Council Discover Project DP200101680. We thank Reid Tingley, Aram Pili and Donald Hobern for their contribution to identifying types of uncertainty in the data.

## Author contributions

The partnership supporting GRIIS includes its development (the IUCN SSC ISSG, GBIF and network of Country Editors), scientific support (M. McGeoch, Q. Groom) and use (CBD, GEO BON, IPBES and Map of Life).

S.P. led the GRIIS Project Team, co-designed the Register, conducted data collection and entry, contacted and communicated with the Country Editors, generated the published checklists and co-wrote the manuscript.

S.B. contributed to data curation, analysis and writing of the manuscript.

P. G. co-designed the Register, and as Chair of the IUCN SSC ISSG led the partnership that resulted in the development of GRIIS.

Q. G, T.H., W.J. and D.S. contributed conceptually to the work and to writing the manuscript. A.R and Y.S. contributed to the analysis and writing of the manuscript.

M.A.M provided technical advice on the design and population of the Register and led the analysis and writing of the manuscript.

**Data Citation**

Pagad, S; Bisset, S; McGeoch, M 2022, Country Compendium of the Global Register of

Introduced and Invasive Species. Dataset. https://doi.org/10.5281/zenodo.6348164

## Footnotes

1 Here, “species” refers to a species, subspecies or lower taxon, and includes any part, gametes, seeds, eggs, or propagules of such species that might survive and subsequently reproduce (sensu https://www.cbd.int/doc/decisions/cop-06-dec-23-en.pdf)

