## Supplementary material for "The Global Register of Introduced and Invasive Species: Country Compendium": Metadata for GRIIS Country Compendium dataset

**GRIIS - Country Compendium Metadata**

Type: CSV (comma delimited)

Encoding: UTF-8

| **Column** | **Is DwC Term?** | **Description** |
| --- | --- | --- |
| recordID | No | Unique identifier for the GRIIS record. |
| scientificName | Yes | The accepted name of the taxon, primarily according to GBIF and in some cases adjusted by GRIIS editors. May contain authorship information. |
| reportedTaxon | No | The scientific name of the taxon as reported by the country or source. |
| species | Yes | The species name of the taxon described in scientificName, excluding subspecies designation and authorship information. Where a genus only record is provided, the suffix "sp." is appended. |
| kingdom | Yes | The kingdom of the taxon described in scientificName. Where unavailable, ‘NOT ASSIGNED’ is used. |
| phylum | Yes | The phylum of the taxon described in scientificName. Where unavailable, ‘NOT ASSIGNED’ is used. |
| class | Yes | The class of the taxon described in scientificName. Where unavailable, ‘NOT ASSIGNED’ is used. |
| order | Yes | The order of the taxon described in scientificName. Where unavailable, ‘NOT ASSIGNED’ is used. |
| family | Yes | The family of the taxon described in scientificName. Where unavailable, ‘NOT ASSIGNED’ is used. |
| taxonRank | Yes | The rank of the taxon described in scientificName.  Terms in field:  GENUS  SPECIES  SUBSPECIES  VARIETY  FORM |
| isHybrid | No | Whether the scientificName is a hybrid name.  Terms in field:  TRUE  FALSE |
| country | Yes | The name of the country according to https://www.un.org/en/about-us/member-states. |
| countryCode | Yes | The alpha-2 code of the country according to the ISO 3166 standard. |
| countryCode_alpha3 | No | The alpha-3 code of the country according to the ISO 3166 standard. |
| habitat | Yes | The environment in which the species occurs or that it is associated with.  Where a species exists in multiple environments, terms are separated with a pipe character (\|).  Terms in field:  TERRESTRIAL  MARINE  BRACKISH  FRESHWATER  HOST |
| establishmentMeans | Yes | The origin (provenance) of the species. ‘Alien’ indicates species that have been introduced outside their natural range by human action. ‘Cryptogenic\|Uncertain’ indicates species whose origins are uncertain. ‘Native\|Alien’ indicates species that are native to one area of the country and alien and invasive in another.  Terms in field:  ALIEN  CRYPTOGENIC\|UNCERTAIN  NATIVE\|ALIEN |
| occurrenceStatus | Yes | Indicates that the taxon is present in the country.  Terms in field:  PRESENT |
| isInvasive | No | Evidence of impact – a systematic decision-making process was used to decide if a species was classified as ‘invasive’ at a country level. See Pagad et al. (2018) for a full description.  Terms in field:  INVASIVE  NULL |
